# Degradation of indole-3-acetic acid by plant-associated microbes

**DOI:** 10.1101/2024.02.08.579438

**Authors:** Lanxiang Wang, Yue Liu, Haoran Ni, Wenlong Zuo, Haimei Shi, Weixin Liao, Hongbin Liu, Yang Bai, Hong Yue, Ancheng Huang, Jonathan Friedman, Tong Si, Yinggao Liu, Mo-Xian Chen, Lei Dai

## Abstract

Plant-associated microbiota affect pant growth and development by regulating plant hormones homeostasis. Indole-3-acetic acid (IAA), a well-known plant hormone, can be produced by various plant-associated bacteria. However, the prevalence of microbes with the capacity to degrade IAA in the rhizosphere has not been systematically studied. In this study, we analyzed the IAA degradation capabilities of bacterial isolates from the roots of Arabidopsis and rice. Using genomics analysis and *in vitro* assays, we found that 21 out of 189 taxonomically diverse bacterial isolates possess the ability to degrade IAA. Through comparative genomics and transcriptomic assays, we identified iac-like or iad-like operon in the genomes of these IAA degraders. Additionally, the regulator of the operon was found to be highly conserved among these strains through protein structure similarity analysis. Some of the IAA degraders could utilize IAA as their sole carbon and energy source. *In planta*, most of the IAA degrading strains mitigated Arabidopsis seedling root growth inhibition (RGI) triggered by exogenous IAA. Importantly, we observed increased colonization preference of IAA degraders from soil to root according to the frequency of the biomarker genes in metagenome-assembled genomes (MAGs) collected from different habitats, suggesting that there is a close association between IAA degraders and IAA-producers. In summary, our findings further the understanding of the functional diversity and roles of plant-associated microbes.

## INTRODUCTION

Indole-3-acetic acid (IAA) is a typical auxin naturally produced by plants, playing a crucial role in various aspects of plant growth and development, such as cell division, elongation, and differentiation [1–4]. IAA is primarily synthesized in developing plant tissues and highly concentrated in the root’s apical part, where the organizing quiescent center accumulates a distinct IAA concentration gradient [5, 6]. Additionally, cells in the root apex exhibit a highly active capacity for IAA synthesis [6]. Alongside root cell exfoliation and other transportation strategies, considerable levels of IAA were detected in root exudates [7–9].

The surface and internal parts of plants are colonized by millions of commensal microorganisms, collectively known as the plant-associated microbiome [10]. Among these habitats, the root is one of the most critical, with numerous studies suggesting that root exudates, such as plant hormones, significantly influence the structure and function of the root-associated microbiome (predominantly the rhizosphere microbiome) [11–13]. Over millions of years of coexistence with their hosts, microbes have evolved multiple strategies for colonization, including the ability to synthesize and catabolize plant-specific metabolites, such as IAA [14]. It is estimated that 80% of commensal microorganisms isolated from the rhizosphere can produce IAA [15]. However, the proportion of the microbes possessing IAA degradation pathways in the rhizosphere is unknown, not to mention the potential ecological roles of these microbes in their habitats.

Two main pathways for IAA consumption among aerobic bacteria have been characterized (Supplementary Figure 1). The gene cluster *iacABCDEFGHIR* (hereafter iac-like operon), responsible for IAA catabolism into catechol, was identified in *Pseudomonas putida* 1290, *Paraburkholderia phytofirmans* PsJN, *Acinetobacter baumannii*, *Enterobacter soli* and *Caballeronia glathei* [16–22] (Supplementary Figure 2). In contrast, the IAA degradation locus *iadABCDEFGHIJKLMNR* (hereafter iad-like operon), resulting in anthranilate as the end-product, was identified in *Variovorax paradoxus* CL14, *Achromobacter* and *Bradyrhizobium japonicum* [23–27] (Supplementary Figure 2). Moreover, among these components, heterologous expression and gene knock-out experimental validations suggested that *iacAE* or *iadDE* are necessary for IAA bio-transformation in *C. glathei* and *V. paradoxus*, respectively [22, 24]. Besides genes encoding enzymes responsible for IAA catabolism, there are components responsible for regulating operon expression in the cluster. The current studies on expression regulation of the operon suggest that most of the iac-like and iad-like operons contain a MarR (multiple antibiotic resistance regulator) family regulator [21, 24, 28]. The crystal structure of iadR and its binding properties were recently determined in *V. paradoxus*, which deciphered the operon expression regulation mechanism [24].

Here, through combining comparative genomics, transcriptomics, with *in vitro* degradation assay, we systematically evaluated the IAA degradation capacity among 189 bacterial strains which were isolated from Arabidopsis (*Arabidopsis thaliana*) and rice (*Oryza sativa*) roots. We predicted the IAA-degrading candidates based on the presence of *iacAE* and *iadDE* in their genomes, followed with experimental validation using the Salkowski method [29] combining with Liquid chromatography-mass spectrometry (LC-MS) analysis. We identified 21 strains belonging to 7 genera, including two previously unreported genera, *Sphingopyxis* and *Curvibacter*, that typically exhibit *bona fide* IAA degradation activity. All IAA degrading strains carry the iac-like or iad-like operon, and the transcriptomic results show that iac and iad gene clusters were upregulated by IAA stimulation. Moreover, the MarR family regulator was present in the operon of all genera with high similarity in their protein structures expect for *Pseudomonas*, which had a putative two-component regulatory system in their iac-like operon. Furthermore, in subsequent assays, we found that some of the IAA degraders could utilize IAA as their sole carbon and energy source. *In planta*, our results demonstrated that exogenous IAA-induced primary root growth inhibition (RGI) was disrupted by most of the IAA degraders, suggesting an important role of IAA degraders in the rhizosphere for host plant growth and development. Finally, by analyzing metagenome-assembled genomes (MAG) and whole genome sequences (WGS) of microbial isolates from different habitats, we found that the prevalence of IAA degraders is positively correlated with naturally occurring IAA resources.

## Results

### Genomic analysis and experimental validation were employed to screen for IAA degraders

Previous reports have identified two types of aerobic auxin catabolic gene clusters in microbes, known as iac-like and iad-like operons [24]. Key genes *iacAE* and *iadDE* play essential roles in IAA degradation [22, 24]. To systematically evaluate IAA degradation capabilities in bacterial commensals isolated from Arabidopsis and rice root, we profiled loci containing genes homologous to *iacAE* and *iadDE* by scanning 189 bacterial genomes from our laboratory collection (Figure 1, Supplementary Table1). A total of 21 strains concurrently containing *iacA* and *iacE*, or *iadD* and *iadE*, were identified as IAA-degrading candidates. Among these, four strains belonging to *Pseudomonas*, five to *Acinetobacter*, and one to *Curvibacter*, show high amino acid sequence identity to the reported iacA and iacE (identity > 60%). Additionally, four strains belonging to *Variovorax* and three to *Achromobacter*, exhibit high amino acid sequence identity when compared to experimentally validated iadD and iadE. A strain belonging to *Sphingopyxis* harbours medium amino acid identity to the iadD and iadE (identity between 40% to 60%). Interestingly, the genomes of three *Sphingomonas* strains contain both iacAE and iadDE with medium amino acid identity (Figure 1), (Supplementary Table2).

**Figure 1.**
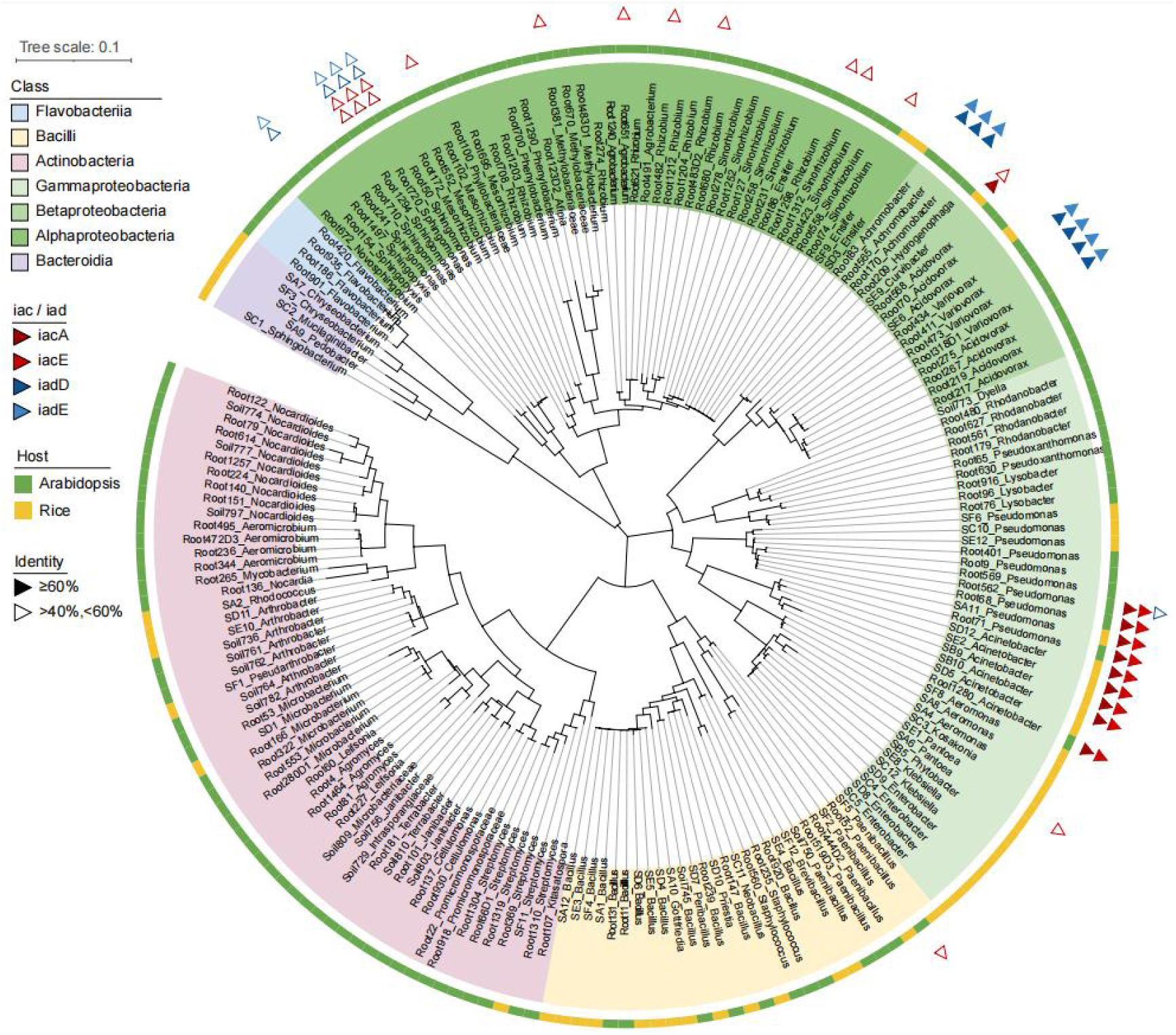
Bacterial strains isolated from Arabidopsis and rice rhizosphere are annotated for genes related to IAA degradation. The 16S rRNA-based phylogenetic tree of the 189 strains was generated with MEGA-X and visualized with iTOL. Strains annotated to have iacA/E or iadD/E (over 40% identity and 60% coverage with template amino acid sequences, see Methods) were labeled with triangles.

To validate the predicted results, we tested the IAA degradation capability among the 137 Arabidopsis root bacterial isolates and 6 IAA-degrading candidates from rice collections by using the Salkowski reaction method. After 72’ hours incubation, bacterial growth and the percentage of IAA degradation of the strains were measured and calculated. A total of 32 out of 143 strains displayed considerable IAA degradation efficiency, such that over 30% of the IAA content were consumed (Supplementary Figure 3A). Among them, 21 IAA-degrading candidates possessing iacAE or iadDE, consumed over 50% of the IAA content in the medium after 72-hour incubation (Supplementary Figure3B). To further detect the consumption of IAA by the IAA-degrading candidates, the supernatant of the culture was analyzed by a highly accurate analytical approach of LC-MS. The specific peak of the IAA was undetectable in these IAA-degrading candidates cultures, suggesting that IAA was degraded by these strains (Supplementary Figure 3C). Ultimately, 21 IAA degraders belonging to seven genera were confirmed through experimental validation, which are consistent with the bioinformatics prediction.

### IAA degraders possess the iac/iad-like operon

To clarify and characterize the IAA catabolism gene clusters in our screened IAA degraders, we performed a BLASTp search using the amino acid sequences of the full iac and iad operons obtained from previous reports against the genomes of 21 strains (Supplementary Figure 2, Supplementary Table 2). A complete iac-like or iad-like operon was identified from the genome of all the IAA degraders except for *Sphingopyxis*_Root154 (Figure 2A). Consistent with previous reports, strains from *Achromobacter* and *Variovorax* possess the iad-like operon, and strains from *Pseudomonas*, *Acinetobacter* and *Sphingomonas* contain the iac-like operon in their genomes. For *Curvibacter_*SE9, and *Sphingopyxis*_Root154, two novel identified IAA-degrading genera, a iac-like operon and a fragmented iad-like operon are present in their genomes, respectively (Figure 2A). Additionally, aside from the complete iac-like operon, two sets of fragmentary iad-like gene clusters were also found in the genomes of strains from *Sphingomonas* (Supplementary Figure 4). Intriguingly, there is another potential fragmented iad-like operon (iad-2) present in the genome of *Sphingopyxis*_Root154, and this operon has the same gene cluster arrangement with strains from *Spingomonas* (Supplementary Figure 4).

**Figure 2.**
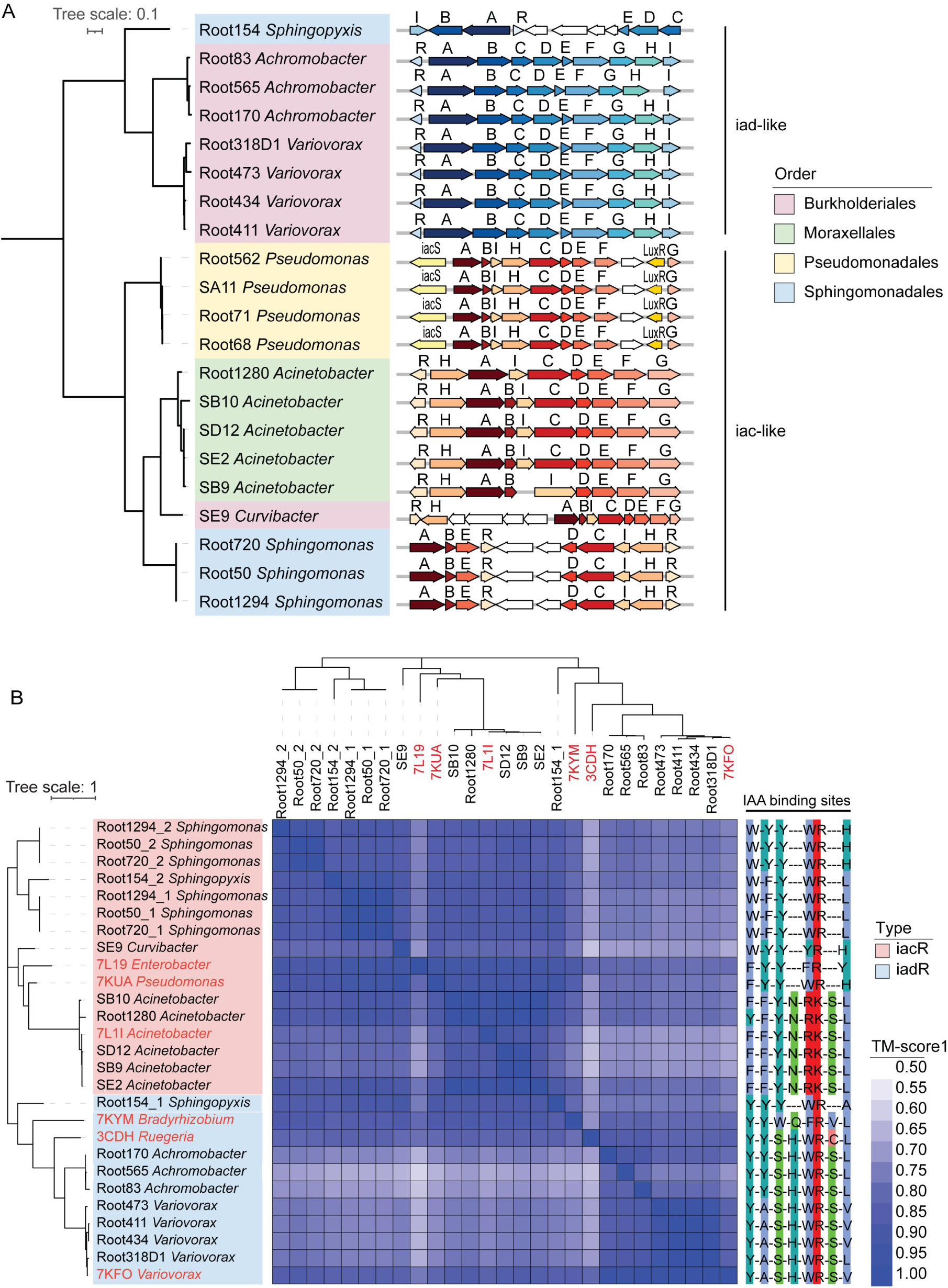
Identification of gene clusters related to IAA degradation and the corresponding MarR family regulators. (A), Organization of *iac* and *iad* gene clusters in 21 IAA-degrading strains. Phylogenetic tree constructed using the concatenated iacA and iacE or iadD and iadE amino acid sequences. Gene transcription directions are indicated by arrows. Letters above the red or blue arrows indicate function genes in the cluster with high protein sequence identity compared to the templates. White arrows indicate genes in this cluster with unknown function. (B), Protein structures of iacR or iadR (all belong to MarR family) are highly conserved among IAA-degrading strains. Heatmap of the protein structural similarity among 27 MarR family proteins displayed the TM1 score. iacR or iadR retrieved from 17 IAA-degrading strains in this study and 6 reference proteins which protein structures and IAA binding sites have been identified. The phylogenetic tree was constructed using the iacR or iadR amino acid sequences. The IAA binding sites of MarR were conservatively distributed in iacR or iadR. Six MarR templates used in this analysis (labeled in red) are 7L1I *Acinetobacter baumannii*, 7KUA *Pseudomonas putida*, 7L19 *Enterobacter soli* ATCC BAA-2102, 7KYM *Bradyrhizobium japonicum*, 3CDH *Ruegeria pomeroyi* DSS-3, 7KFO *Variovorax paradoxus* CL14.

To investigate the evolutionary relationships among different IAA degraders, we constructed a phylogenetic tree using concatenated iacAE or iadDE amino acid sequences. Intriguingly, strain organization in this tree differed substantially from the phylogenetic relationships based on 16S rRNA sequences (Supplementary Figure 5). *Curvibacter* was more closely related to *Acinetobacter* than to other Burkholderiales. *Sphingomonas* and *Sphingopyxis*, both belonging to Sphingomonadales, appeared on different branches. Moreover, *Sphingomonas* genomes contained iac-like operon, while the *Sphingopyxis* genome harbored an iad-like operon (Figure 2A). These results suggest that, evolution of microbes un-synchronized with some gene cluster acquisition.

### The MarR family regulators exhibit high degree of structural conservation among IAA-degrading strains

In all IAA-degraders except *Pseudomonas*, both iac-like and iad-like operons contain one or two MarR-family regulator (R) that are involved in the operon expression regulation (Figure 2A). In contrast to previous reports [24], our four *Pseudomonas* strains exhibited the involvement of a putative two-component regulatory system in their iac-like operons (Figure 2A). Upon comparison with the iac operon in the related reference strain *Pseudomonas putida* 1290, it seems that the components of the cluster may have been acquired from a distant genus, such as *Paraburkholderia phytofirmans* (Supplementary Figure 2).

Generally, the MarR regulator acts as a negative transcriptional factor for the operon since it binds to the upstream DNA region, inhibiting operon expression. Due to the similar role of this protein in regulating iac and iad operon expression, we hypothesized that they are homologous with high amino acid sequence identity. To verify our hypothesis, multiple protein sequence alignment was performed using 21 potential MarRs identified from our 17 strains and six MarR reference proteins [24]. Consistent with previous reports, MarR regulators are diverse, as they have low protein sequence identity (Supplementary Figure 6). The phylogenetic tree based on the protein sequence of 27 MarRs demonstrated that iacR naturally separated from iadR (Figure 2B), which is consistent with previous reports [30]. MarRs are relatively conserved within a genus, while iacRs display more sequence diversity. It is worth noting that three strains from *Sphingomonas* contain two iacRs in their iac operons, respectively (Figure 2 and Supplementary Figure 6). Also, two MarRs were screened from Root154_*Spingopyxis* genome with low similarity, Root154_1 (gene_00256) and Root154_2 (gene_00334) (Figure 2B, Supplementary Figure 6). In addition, except for Root154_1 (gene_00256), other MarRs from Sphingomonadaceae were grouped together and classified as iacR (Figure 2B).

To further elucidate the mechanism of functional conservation among MarRs, the predicted protein structures of 21 hypothetical MarRs were generated using AlphaFold2 [31] (Supplementary Figure 7). The protein structures of six referential MarRs were retrieved from the PDB, and pairwise comparisons were performed among the 27 MarRs using TM-align [32]. As shown in the heatmap, all pairwise comparisons obtained a high TM score (all pairwise TM score > 0.5) [33] (Figure 2B), suggesting that all MarRs displayed high similarity in their protein structures. In addition, the IAA binding sites of MarRs also exhibited high conservation, especially within a genus level (Figure 2B).

### Iac-and iad-like operons were up-regulated by IAA

To explore how iac-or iad-like operons were regulated by IAA, we examined the transcriptomes of five strains in response to this compound. The strains selected for RNA-seq were *Pseudomonas*_Root71, *Achromobacter*_Root170, *Variovorax*_Root473, *Curvibacter*_SE9, and *Sphingopyxis*_Root154. They were cultured with M9 minimal medium supplemented with IAA or glucose, and cells were collected in specific time point (see method). Among them, *Curvibacter* and *Sphingopyxis* were studied for IAA degradation activity for the first time. In total, 308, 151, 274, 871, and 220 differentially expressed genes (DEGs) were identified in these five strains, respectively (Supplementary Table 3). Both iac-like and iad-like operon were significantly upregulated by IAA, except for Root154 (Figure 3). Furthermore, categorization of up-and down-regulated DEGs indicated that IAA treatment broadly influenced many metabolism pathways (Supplementary Figure 8).

**Figure 3.**
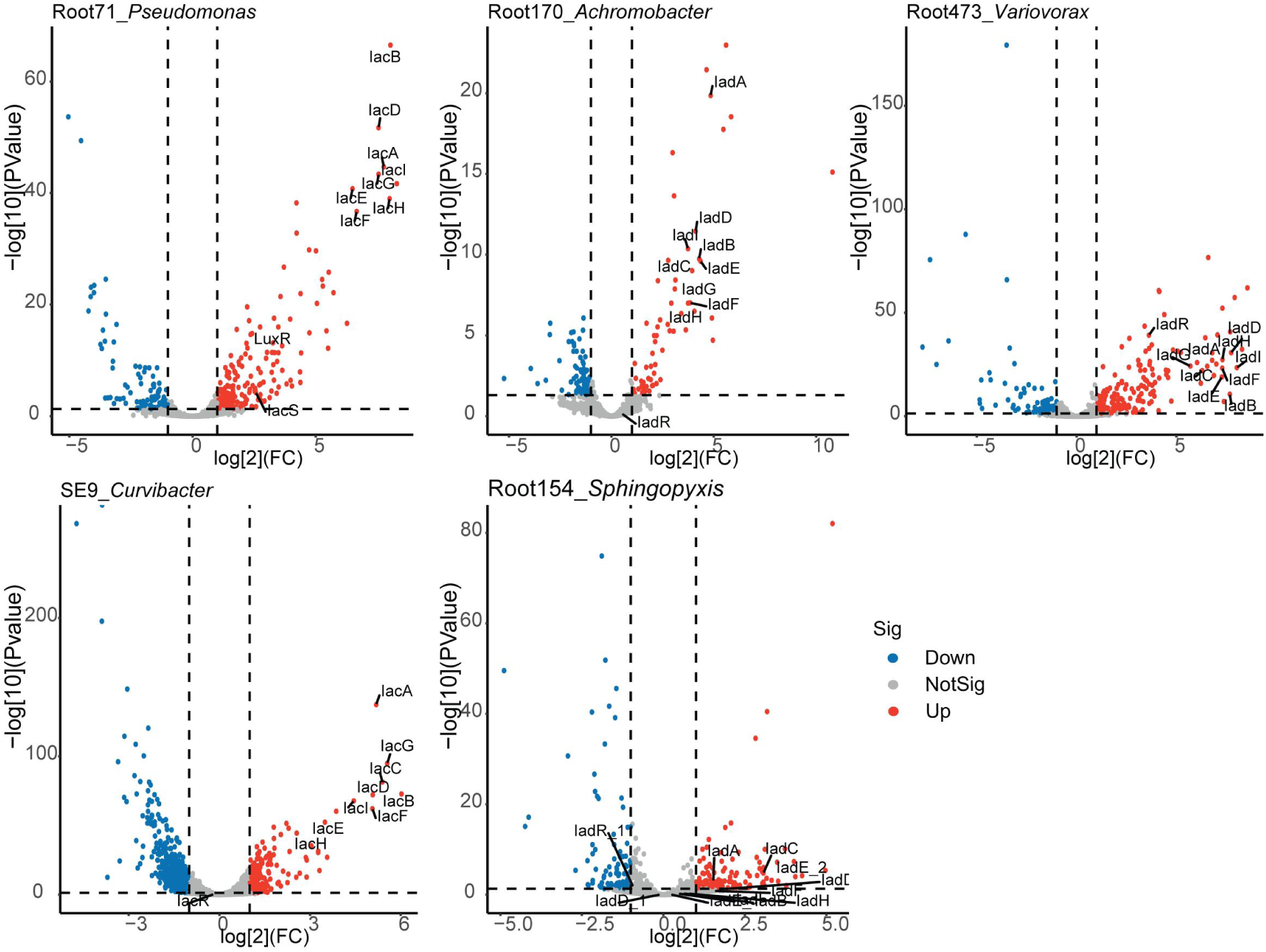
RNA-seq reveals that the transcription of the iac-like or iad-like gene cluster is induced by IAA. The 5 selected strains for RNA-seq analysis were cultured with M9 medium supplemented with either IAA or glucose as the sole carbon source. The deferential expression genes between IAA and Glucose treatments were analyzed. Significantly up-regulated or down-regulated genes were identified with log ^FoldChange^ ≥ 1 or ≤ -1 and adjusted *p*-value < 0.05 as cutoffs. Three individual colony of each strain were used to perform the transcriptome analyses.

It has been reported that chemicals in root exudates may enhance plant-microbe interactions through transcriptional regulation of bacterial motility [34]. Flagella are organelles used by microbes for movement. Notably, in Root473 and Root154, genes involved in flagellar biosynthesis, and chemotaxis proteins were upregulated by IAA treatment when compared with glucose control, suggesting that for these two strains, IAA might be more attractive than glucose. However, in SE9, genes encoding flagellum-related DEGs and chemotaxis proteins were down-regulated by IAA stimulation, indicating that IAA elicited different molecular responses in different strains. Consistent with previous reports, catechol, the end-product of iac-pathway, will be further catalyzed by downstream enzymes, catABC [16, 20, 21]. Genes involved in catechol pathway in Root71 and Root170 were up-regulated by IAA treatment (Supplementary Table 3, highlighted).

### IAA can be utilized as the carbon source

To evaluate the utilization of IAA by the IAA degraders, *in vitro* assay were preformed among the screened 21 IAA degraders. The IAA degradation efficiency and bacterial growth of the strains were carried out in M9 minimal medium with exogenous IAA. Results showed that except for strains from *Sphingomonas* and *Sphingopyxis*, other strains benefited from IAA as the sole carbon source and completely consumed IAA within 72 hours (Figure 4A). Strains from *Acinetobacter* presented the maximum degradation and growth rates, consuming IAA completely within 12 hours (Figure 4A and B). Compared with strains from *Acinetobacter*, strains from *Pseudomonas* displayed the equivalent IAA degradation efficiency and less biomass. Strains from *Variovorax*, *Achromobacter*, and *Curvibacter*, which generally degraded IAA completely within 60 hours, also showed extremely slow growth rate may suggest that cell proliferation of these strains require more carbon and energy source. On the other hand, strains from *Sphingomonas* only consumed partial IAA in the M9 minimal medium, while *Sphingopyxis* barely utilized this compound under this condition of culture. Combined with the transcriptome analysis, our results here suggest that IAA triggered the expression of iac/iad operon and biotransformed this compound in the medium, although the by-products of IAA degradation may not suitable for cell growth as sole carbon and energy source.

**Figure 4.**
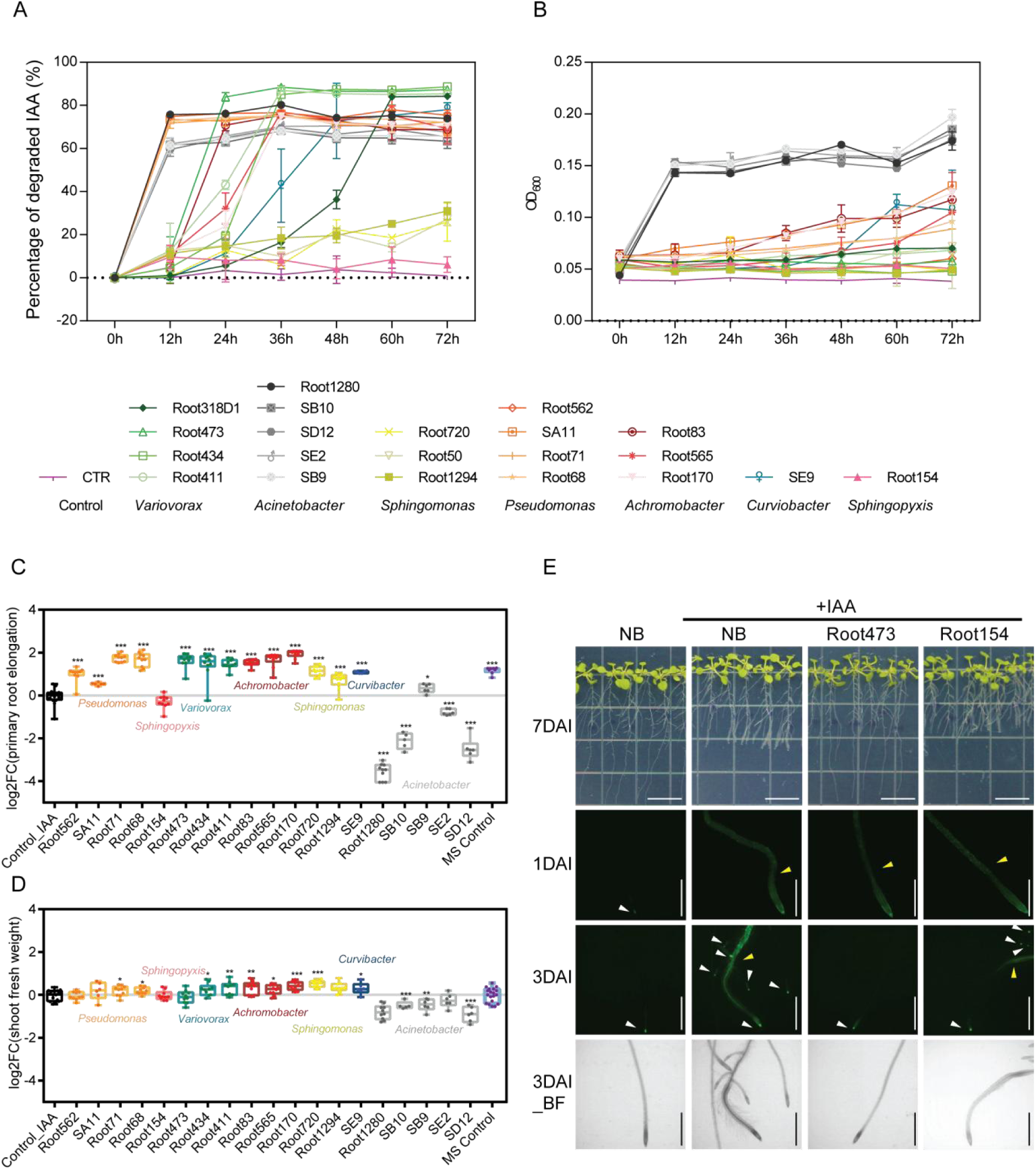
IAA-degrading strains can utilize IAA and suppress the IAA-induced root growth inhibition. (A), IAA consumed by IAA degraders *in vitro* and (B) their growth as measured by OD_600_ in M9 minimal medium supplemented with IAA as the sole carbon source (n=4). (C), Except for strains from genus of *Sphingopyxis* and *Acinetobacter*, IAA-induced root growth inhibition was suppressed by IAA-degrading strains. Boxplot middle line, median log2 fold change of primary root elongation (vs Control_IAA); box edges, 25th and 75th percentiles; whiskers, 1.5× the interquartile range. Left to right n = 16, 10, 6, 10, 10, 10, 10, 10, 10, 10, 10, 10, 10, 10, 6, 10, 5, 6, 6, and 8 biological replicates. (D), Apart from *Acinetobacter*, strains inoculation have no negative effects on shoot fresh weight. Boxplot middle line, median log2 fold change of shoot fresh weight (vs Control_IAA); box edges, 25th and 75th percentiles; whiskers, 1.5× the interquartile range. Left to right n = 17, 10, 10, 6, 10, 10, 10, 10, 10, 10, 10, 10, 10, 10, 10, 10, 7, 9, 9, and 19 biological replicates. (E), Images of representative Col-0 seedlings grown axenically (NB) or with IAA-degrading strain inoculation. Upper panes show the representative images of seedlings grown on 1/2 MS agar plate supplemented with 100nM IAA at 7 days after inoculation with or without strain. Bar = 1.4cm. The other planes show the representative primary root images of *DR5::GFP* plants after inoculated with strain for 1 and 3 days. Bar = 1mm. Write arrows show the GFP signal on root tips. Yellow arrows show the GFP signals on root which were induced by exogenous IAA. BF, bright field. Significant differences compared with control group were determined using Student’s *t*-test: \**P*<0.05, \*\**P*<0.01, \*\*\**P*<0.001.

### IAA degraders contribute to the regulation of plant root growth

Auxin homeostasis in plant roots is achieved through local synthesis, polar transport, and the contribution of IAA-producing/consuming microorganisms, which is crucial for root growth. To clarify the biological role of the root isolates possessing IAA degradation capability identified in this study, seven-day-old Arabidopsis seedlings were transferred to 1/2 MS agar medium supplemented with 100nM IAA and inoculated with the IAA degraders individually. After additional seven days of inoculation, primary root elongation was measured. Primary root elongation was suppressed when exogenous IAA was added to the medium (Figure 4C and E). In mono-associations, normal primary root growth was observed when seedlings were inoculated with strains from *Pseudomonas*, *Variovorax*, *Achromobacter*, *Sphingomonas*, *Curvibacter*, and SB9, the only strain from *Acinetobacter* (hereafter referred to as RGI-suppressive IAA degraders, while the rest are RGI-non-suppressive IAA degraders) (Figure 4C and E, Supplementary Figure 9). Consistently, a small range of fresh weight enhancement of shoots was observed in these positive mono-associations (Figure 4D). On the other hand, the rest strains from *Acinetobacter* inhibited primary root elongation, as well as shoot fresh weight (Figure 4C and D). No significant effects on root growth or shoot fresh weight were observed in *Sphingopyxis* treatment.

To further explore whether the restoration of RGI by RGI-suppressive IAA degraders is directly related to auxin signaling in plants, the Arabidopsis auxin reporter line *DR5::GFP* was treated with IAA and simultaneously inoculated with IAA degraders. Fluorescence of the *DR5::GFP* induced by exogenous IAA remained stable in axenic control at day 1 and day 3 post-inoculation. The GFP signal in *DR5::GFP* roots was quenched at day 3 after inoculation with RGI-suppressive IAA degraders. Consistent with the root elongation phenotype, RGI-non-suppressive IAA degraders, including strains from *Sphingopyxis* and *Acinetobacter*, could not quench the root fluorescence caused by exogenous IAA (Figure 4E, Supplementary Figure 9). Interestingly, SB9, the RGI-suppressive IAA degrader from *Acinetobacter*, quenched the fluorescence at day 3 (Supplementary Figure 9). To rule out the possibility that the phenomenon observed in RGI-non-suppressive IAA degraders was not caused by failed colonization, colonization of strains was investigated by calculating colony-forming units (CFUs), which were further normalized to root fresh weight. After seven days of inoculation, all strains successfully colonized roots, and exogenous IAA supplementation had no significant effect on bacterial colonization (Supplementary Figure 10 and 11).

### Catalogue of potential IAA-degrading bacteria from diverse habitats

IAA degrading bacteria in the rhizosphere play an important role in maintaining auxin homeostasis in roots to ensure normal plant growth and development [25]. Nevertheless, their distribution across various habitats remains limited. To ascertain the distribution of IAA-degrading strains in different habitats, we analyzed a large-scale survey of 11,586 high-quality MAGs (including 750 isolates) collected from mammal gut, aquatic environment, soil and plants (Figure 5) (Supplementary Table 4) [35–43]. The iacA/E and iadD/E were used as the biomarkers to screen the genomes of the collections. We noted that no hits were identified in MAGs collected from cold seeps or animal/human guts, which are reasonable since these habitats are normally hypoxic, while the iac/iad pathways are aerobic (Supplementary Table 4). Furthermore, iac or iad-like operon is also absent in the facultative anaerobic environment of the human oral [43]. In contrast, among the 692 MAGs collected from human skin, five potential IAA degraders containing iac-like operon were identified [43].

**Figure 5.**
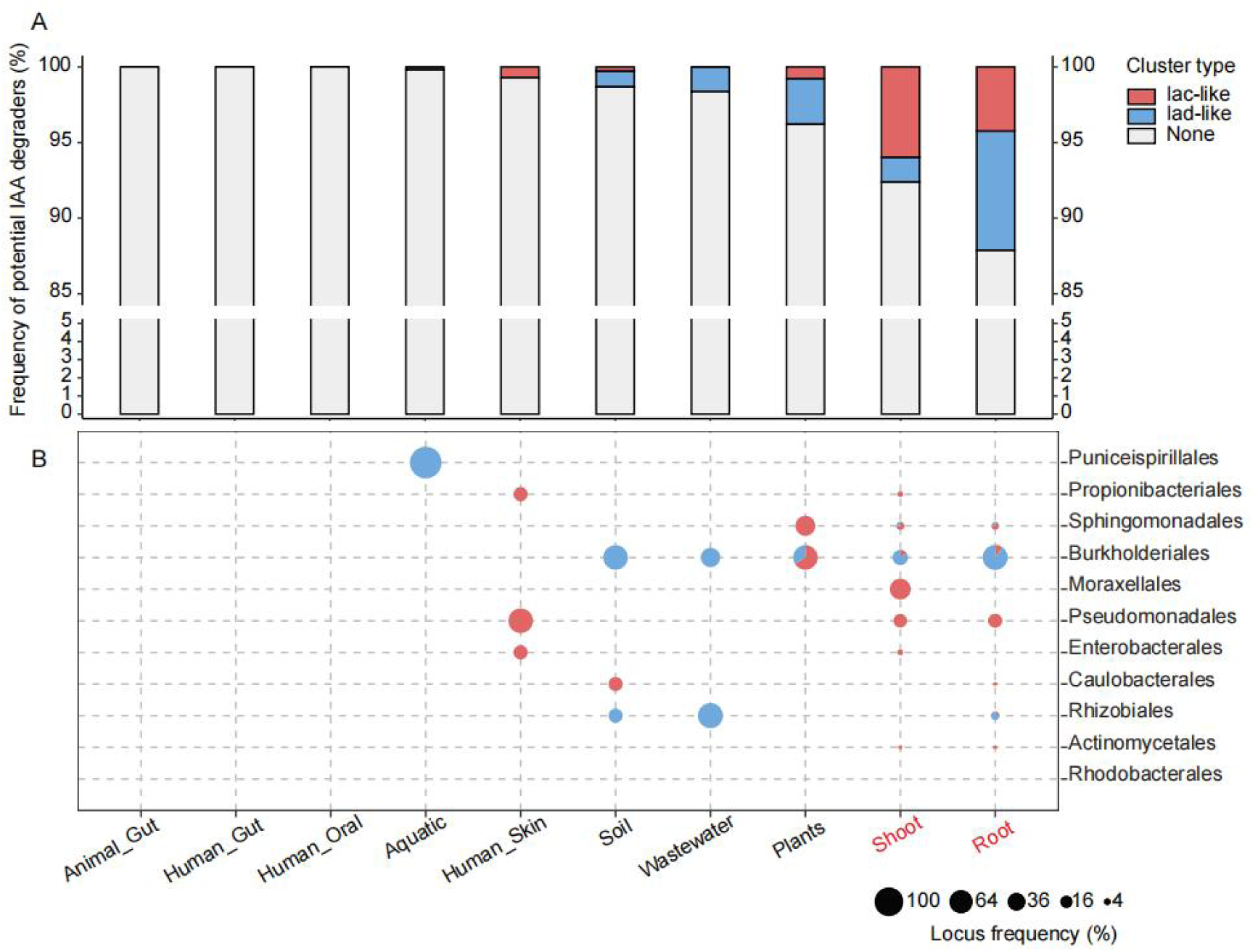
The distribution of IAA-degrading bacterial strains across various habitats. (A), The frequency of potential IAA degraders whose genome contains iacA and iacE or iadD and iadE in different habitats. IAA degradation types were labeled with red or blue color. Ggbreak was applied in this analysis [65]. (B), IAA degradation types are varied at bacterial order level. In total, 11,586 high quality MAGs and isolates were analyzed in this study. Samples containing isolates were highlighted with red color. All MAGs were ≥90% complete, were ≤5% contaminated and had a quality score (completeness -5 × contamination) of ≥65.

On the other hand, more potential IAA degraders were identified from environment samples, especially plant-associated samples. 0.65% MAGs (2 out of 304) collected from marine samples [43], and 1.63% MAGs (8 out of 492) collected from wastewater samples were identified harboring iad-like operon [42]. In soil samples, 1.31% MAGs (5 out of 382) were identified containing iac or iad-like operon [35, 42]. Within the plant collections, 3.79% MAGs (5 out of 132) affiliated with Sphingomonadales and Burkholderiaceae were identified as containing the iac or iad-like operon [42]. Furthermore, 7.60% MAGs (60 out of 789, containing 206 isolates) collected from plant shoot [36, 38, 40] and 12.13% MAGs (66 out of 544, all are isolates) isolated from plant root [40, 41] were found to harbor the IAA degradation operon. Overall, there is a consistent increase in the frequency of potential IAA degraders from aquatic to terrestrial environments, from soil to plants, and from plant shoots to roots. Moreover, potential IAA degraders containing iac-or iad-like operon and belonging to Burkholderiales were widely distributed across various habitats (Figure 5).

## DISCUSSION

In this study, combining genomic analysis and experimental validation, we performed a systematic screening of IAA degraders isolated from Arabidopsis and rice root. We found that 21 strains belonging to 7 genera exhibit remarkable IAA degradation activity. In addition to the previously reported *Pseudomonas*, *Achromobacter*, *Variovorax*, *Acinetobacter*, and *Sphingomonas*, strains from *Sphingopyxis* and *Curvibacter* also displayed outstanding IAA-degrading activity. The genomes of all IAA degraders contain iac-or iad-like IAA degradation operon, and these operons were upregulated by IAA treatment. By integrating protein sequence alignment with protein structural similarity analysis, we revealed that the MarR family regulators are structurally conserved within genus. *In vitro* assays suggested that the screened 21 strains are *bona fide* IAA degraders. However, only a subset of these strains directly depleted exogenous IAA to maintain the host-plant root growth. Intriguingly, MAGs analysis showed that IAA-degrading candidates naturally colonized plant-associated habitats, which aligns with the notion that IAA degraders are prevalent in habitats closely linked to IAA producers. Our findings revealed a key role of IAA degraders inhabiting in plant and the underlying degradation mechanism.

Comparative genomics studies indicate that the gene clusters of IAA degraders were probably acquired in a natural environment through a horizontal gene transfer pathway, with selective loss, duplication, and rearrangement of IAA-degrading genes during evolution. The gene cluster contains structural genes encoding enzymes responsible for IAA degradation and a set of genes involved in gene expression regulation and compound transportation, and the core components are highly conserved. The structure and arrangement of the iad-like operon in *Variovorax* are highly similar to that in *Achromobacter*, indicating that these two genera belonging to Burkholderiales may have obtained the operon from the same ancestor/donor. In contrast, *Curvibacter*, another genus belonging to Burkholderiales, possesses the iac-like operon, suggesting the possibility that IAA degradation gene clusters were obtained through horizontal gene transfer (Supplementary Figure 5). *Sphingomonas* and *Sphingopyxis* are closely related and both belong to the Sphingomonadaceae. A complete iac-like operon uniquely present in the genomes of strains from *Sphingomonas*. Moreover, IAA-degrading gene cluster mining analysis showed that two additionally fragmentary iad-like operons exist on *Sphingomonas* genomes, and one of them has the same gene arrangement to the *Sphingopyxis* (Supplementary Figure 4), suggesting the evolutionary homology of these two genera. Additionally, although *Acinetobacter* and *Curvibacter* belong to different families, both genera have iac-like operons and the operons have the similar gene arrangement, suggesting that they may obtained the gene cluster from a closer donor (Figure 2A).

IacR and iadR are an essential component of the IAA-degrading gene cluster, and functional studies suggest that it normally serves as an expression suppressor of the operon. Recently, the protein crystal structure of iadR was resolved, and it was confirmed that iadR binds to the upstream DNA sequence of *iadA* to inhibit iad locus expression in *Variovorax paradoxus* CL14 [24]. The presence of IAA results in iadR being released from the DNA binding site, further disinhibits the expression of the iad operon. The MarR regulators have highly conserved protein structures despite having highly differentiated protein sequences (Supplementary Figure 6 and 7, Figure 2B). This may explain why different MarR regulators perform similar functions in regulating operon expression. Similar to *Paraburkholderia phytofirmans* PsJN, LuxR and iacS uniquely exist in the iac operons of four strains belonging to *Pseudomonas*, suggested that a putative two-component regulatory system independently evolved or obtained in this genus (Figure 2A) [20].

Plant growth requires multifaceted regulation, including but not limited to IAA production and degradation by plant, as well as auxin regulation by plant-associated microbiota [25]. It is estimated that 80% of rhizosphere commensal isolates possess the capability of producing IAA [15]. While, based on our results, only 11.11% of the isolates (21 out of 189 in this study) and 7.84% (131 out of 1465) of the plant-associated MAGs (Supplementary Table 4) exhibit potential IAA-degrading capability. The results of mono-association further reveal the biological role of IAA degraders in manipulating IAA homeostasis in plants to maintain root growth. However, not all IAA-degrading strains have the ability to reverse the severe inhibition of root growth induced by excess IAA. Moreover, our mono-association results showed that *Acinetobacter* has negative effects on plant growth (Figure 4, Supplementary Figure 9), which is inconsistent with recent reports that *Acinetobacter* can act as a plant growth-promoting rhizobacteria (PGPR) [44, 45]. In addition to being a strong IAA-degrading bacterium, *Acinetobacter* is also one of many that produce IAA and can thus sabotage plant physiology by adding to the endogenous IAA pool in plants [46]. Given these contradictory traits, the biological function of *Acinetobacter* for plants and the underlying mechanism need to be further studied.

Our large-scale MAGs analysis results suggested that the prevalence of IAA-degrading taxonomy increased gradually from aquatic to terrestrial environments, from soil to plants, and from plant shoots to roots (Figure 5). It is reasonable that IAA-degrading bacteria may be recruited to utilize IAA as sources of carbon and/or nitrogen, whereas plants and numerous plant-associated IAA-producing microbes serve as natural sources of this compound.

Lastly, hormones may play a crucial role in mediating the interactions between hosts and microbes. For instance, *Mycobacterium neoaurum* possessing the capability to degrade testosterone was isolated from the fecal samples of testosterone-deficient patients with impression. Further experiments revealed the potential association between human gut microbes expressing 3β-HSD and depressive symptoms resulting from testosterone degradation [47]. Beyond IAA, other plant hormones have been reported to be synthesized or metabolized by microbes. For examples, rhizobacteria such as *Rhodococcus* sp. P1Y and *Novosphingobium* sp. P6W have been reported to utilize abscisic acid (ABA), consequently stimulating plant growth through an ABA-dependent mechanism [48]. We envision that investigations on microbial metabolism of host hormones may offer novel insights into the understanding of homeostasis, host physiology and the development of diseases.

## MATERIALS AND METHODS

### Plant materials and bacterial strains

*Arabidopsis thaliana* ecotype Columbia (Col-0) was obtained from laboratory stock. The Arabidopsis auxin reporter transgenic line *DR5::GFP* was kindly provided by Prof. Xugang Li (Shandong Agriculture University). The 137 bacterial commensals isolated from Arabidopsis roots or soil were gifts from Prof. Paul Schulze-Lefert (Max Planck Institute for Plant Breeding Research, Cologne, Germany)[40]. Detailed information on individual strains can be found at At-RSPHERE (http://www.at-sphere.com/).

The 52 rice root-associated bacterial isolates analyzed in this study were retrieved from our laboratory stock. In detail, rice root samples contain bulk soil were collected in field and immediately delivered with ice to our lab. After removed the bulk soil, 10g root samples contain rhizosphere soil were washed multiple times with sterilized water until there is no obvious soil on root surface. The washings were mixed as the rhizosphere sample. Rice roots were then grind with 10ml 1 × PBS buffer in a sterilized mortar and filtered with sterilized gauze. Both samples were then spread on the surface of 1/5TSB agar plates with series dilution. After incubation for 3 days at 25℃, single colonies were randomly picked from the plates for twice purification. A total of 207 isolates were identified at the species level by sequencing 16S rRNA gene with the primers 27F (AGAGTTTGATCCTGGCTCAG) and 1492R (GGTTACCTTGTTACGACTT). For whole genome sequencing, the genome DNA of selected 72 strains were individually extracted using FastDNA Spin Kit for Soil (MP Biomedicals, USA). Library preparation was performed using the Hieff NGS OnePot II DNA Library Prep Kit (Yeasen, China) for Illumina with 50 ng DNA per sample. The draft genomes were generated with the HiSeq Xten platform (Illumine, USA). Quality control of the raw reads were filtered with fastp [49], followed with genome assembly through Unicycler[50]. CheckM was used to estimate the quality of each genome, including the numbers and N50 of the contigs, the contamination, and the completeness[51]. Prokka was used to annotate the function of all assembled genomes[52]. Taxonomy annotation of the isolates was performed using the Genome Taxonomy Database Toolkit (GTDB-Tk)[53] with reference to GTDB release 207[54]. Isolates were assigned at the species level if the ANI to the closest GTDB-Tk representative genome was ≥95% and the aligned fraction was ≥60%. General taxonomical information of these 189 strains is listed in Supplementary Table 1.

### Construction and modification of phylogenetic tree

The phylogenetic trees were constructed using MUSCLE[55] for multiple sequence alignment and MEGA[56] for tree construction. Specifically, Figure 1 was constructed using the 16S rRNA from 189 strains; Figure 2A was constructed using the amino acid sequences of key genes involved in IAA metabolism from 21 IAA-degraders; Figure 2B was constructed using the amino acid sequences of 21 MarR proteins discovered from the 17 IAA-degraders and 6 related templates. The generated phylogenetic trees were further visually modified using iTOL[57].

### Identification of potential IAA-degraders

Prokka[52] was used for functional annotation of isolates and MAGs derived from different environments, meeting the criteria of genome completeness (≥90%) and contamination (≤5%). Diamond[58] was employed for sequence alignment of annotated genomes, using previously reported IacA and IacE or IadD and IadE sequences as templates. The alignment filtering threshold was set at sequence similarity (Identity) >50% and sequence coverage (Coverage) >60%. If both gene combinations were simultaneously identified in a genome and determined to be located in the same gene cluster (IadD adjacent to IadE; IacA with a distance less than 7 Coding Sequences (CDS) from IacE), the strain was considered to possess IAA degradation capability.

### Bacterial culture and screening of IAA degradation

Individual bacteria from glycerol stock were incubated on 1/2 tryptic soy broth (TSB, Sigma-Aldrich, USA) agar plates at 25°C for 5 days. A single colony of each strain was then cultured in 1/2 TSB liquid medium at 25°C with 400 rpm shaking. When the cell culture reached the exponential growth phase, the optical density of the culture was measured at 600 nm (OD_600_) using a SynergyTM H1 microplate reader (BioTek, USA).

The bacterial culture was washed once with 1×PBS and then added to 1 mL of 1/2 TSB medium supplemented with or without 0.4 mM IAA (Sigma-Aldrich, USA) to achieve a final OD_600_ of 0.05. After 72 hours of incubation at 25°C with 400 rpm shaking, the IAA content of each sample was measured using the Salkowski method [29]. Briefly, 120 μL of Salkowski reagent was mixed with 60 μL supernatant of culture, and the absorbance of the mixture was measured at 530 nm after 30 minutes of incubation in the dark. The remaining IAA contents in the medium were then calculated using IAA standard curves. The bacterial degradation rate was further calculated as the IAA consumed divided by the initial IAA content in the culture.

### Validation of IAA degradation by LC-MS analysis

To validate the IAA degradation results of the Salkowski method, cell cultures of selected strains were further analyzed by LC-MS [29]. After 72 hours of incubation with IAA, the cell cultures were collected to test the content of IAA in the medium. In detail, the residual IAA of the cell cultures was extracted twice with ethyl acetate, followed by volatilization, and the residue was further dissolved in 80% methanol. After filtration, a 2 μL sample was separated using a C18 column (Infinity Lab Poroshell 120 EC-C18, 2.1 × 50 mm, 2.7 μm; Agilent, USA) connected to the Agilent 6470B triple quadrupole LC/MS (Agilent, USA). Solvent A (water supplemented with 0.1% formic acid) and solvent B (acetonitrile supplemented with 0.1% formic acid) were used as mobile phases at a flow rate of 0.2 mL/min under a gradient elution: 0-2 min, 10% B; 2-8 min, 40% B; 8-11 min, 70% B; 11-15 min, 10% B. The quantification of IAA extracted from culture and IAA standards was performed using the positive-ion multiple reaction monitoring (MRM) method.

### Growth experiment with IAA as the sole carbon source

Selected IAA degraders were individually cultured at 25°C with 400 rpm shaking in 1 mL M9 minimal salts medium (Sigma-Aldrich, USA), supplemented with 2 mM MgSO_4_ and 0.1 mM CaCl_2_. A 0.4 mM IAA solution was added to the culture as the sole carbon source. OD_600_ and IAA concentrations were measured at 7 time points: 0 h, 12 h, 24 h, 36 h, 48h, 60 h and 72h.

### RNA-seq and data analysis

*Pseudomonas*_Root71, *Achromobacter*_Root170, *Variovorax*_Root473, *Curvibacter*_SE9, and *Sphingopyxis*_Root154 were grown in 1 mL M9 medium supplemented with 1.712 mM glucose or 1 mM IAA. The initial OD_600_ of the cultures was 0.05, and they were incubated at 25°C with 400 rpm shaking. Cells from *Pseudomonas*_Root71, *Achromobacter*_Root170, *Variovorax*_Root473, *Curvibacter*_SE9, and *Sphingopyxis*_Root154 were collected for RNA-seq at 14 h, 16 h, 14 h, 48 h, and 20 h, respectively. Bacterial pellets were collected by centrifuging the culture at 12,000 rpm for 10 min at 4°C. The pellets were stored at -80°C until RNA extraction.

Further experiments including RNA extraction, library preparation and sequencing were preformed at Magigene Co. Ltd using the Nova Seq6000 platform (Illumine, USA). In detail, bacterial RNA samples were extracted using the Trizol followed with quality control by Thermo NanoDrop One and Agilent 4200 Tape Station. Epicentre Ribo-Zero rRNA Removal Kit was using to remove Ribosome RNA in the samples. Library preparation was performed using the NEBNext Ultra Directional RNA Library Prep Kit for Illumina (New England Biolabs; USA) with 1 μg total RNA per sample. The raw data were processed with RNA-seq pipeline from nf-core (nf-core/rnaseq, v3.11.1) [59], with trimming enabled. The clean reads were then mapped to the reference genome of corresponding strain with STAR (v2.7.10a) [60]. Gene quantification was subsequently done using Salmon (v1.5.2) [61]. Differentially expressed genes (DEGs) were identified using DESeq2 (v1.36.0) [62] with log2FoldChange ≥ 1 or ≤ -1 and adjusted *p*-value < 0.05 as cutoffs. Three biological replicates of each sample were used to perform the transcriptome analyses.

### Plant experiments

Strains selected for *in planta* assays were pre-cultured in 1/2 TSB at 25°C for 2-3 days until cloudy. On the day of inoculation, the bacterial culture was subcultured at a 1:3 ratio for an additional 5 hours. A 500 μL aliquot of bacterial culture was centrifuged at 6,500 × g for 2 min. After washing twice with 1 × PBS buffer, the bacterial pellets were resuspended in 1×PBS buffer and adjusted to an OD_600_ of 0.01. A 100 μL aliquot of bacterial suspension was then spread on half-strength Murashige and Skoog (MS) (Sigma-Aldrich, USA) plates supplemented with or without 100 nM IAA.

Arabidopsis seeds were surface-sterilized with 75% ethanol for 1 min, 20% bleach for 15 min, and rinsed 5 times with sterile distilled water. Seeds were sown evenly on 1/2 MS plates with 0.5% agar and 3% sucrose. After 2 days of stratification at 4°C in the dark, seeds were vertically grown in a growth chamber under a 16-h dark/8-h light regime at 22°C for 7 days. Ten seedlings were transferred to the prepared half-strength MS plates containing IAA and IAA-degrading bacteria. The initial position of the root tip was labeled, and after an additional 7 days of growth, the final position of the root tip was labeled again. Pictures of the plates were captured with a camera (Nikon, Japan), and the elongation of the primary root was measured using ImageJ [63].

### Fluorescence microscopy

GFP fluorescence in the roots of *DR5::GFP* transgenic lines was visualized using a Ti2-E fluorescence microscope (Nikon, Japan) at 1 and 3 days after inoculation, respectively. The experiment was performed in two independent replicates.

### Measurement of bacteria root colonization

Colony-forming units (CFUs) were counted as previously described with minor modifications [64]. Briefly, after 7 days of inoculation, roots were separated from shoots using a sterile scalpel, taking care to avoid contamination between different bacterial treatments. Two roots were placed in pre-weighed sterile tubes containing metal beads, and the tubes were weighed again to obtain the fresh weight of the roots. Root samples were then homogenized using a TissueLyzer (Shanghai Cebo, China) at 30 Hz for 30 seconds. A 500 μL aliquot of 1 × PBS buffer was added to the tube, and the samples were serially diluted in a sterile 96-well plate. A 5 μL sample was dropped onto a 1/2 TSB plate, and the plate was flipped sideways to allow the liquid to flow evenly. Finally, the plates were placed at 25°C for two days until single colonies appeared. The colonization ability of each strain on the root was calculated according to the CFU count.

## ACKNOWLEDGEMENTS

We gratefully acknowledge Prof. Paul Schulze-Lefert for sharing Arabidopsis associated microbes. We gratefully acknowledge Prof. Xugang Li for providing Arabidopsis auxin reporter transgenic line *DR5::GFP*.

The authors acknowledge the financial support from the National Natural Science Foundation of China (No.32061143023), Guangdong Basic and Applied Basic Research Foundation (No.2023A1515012006), Platform funding for Guangdong Provincial Enterprise Key Laboratory of Seed and Seedling Health Management Technology (2021B1212050011), Guizhou Provincial Basic Research Program (Natural Science)-ZK[2023]-099, the Program of Introducing Talent to Chinese Universities (111 Program, D20023), the Frontiers Science Center for Asymmetric Synthesis and Medicinal Molecules, Department of Education, Guizhou Province [Qianjiaohe KY (2020)004].

## AUTHOR CONTRIBUTION

LD and MXC designed and supervised the project. LXW and YL conducted the experiments. HRN, WLZ and HBL performed the bioinformatics analysis. HMS and WXL isolated the rice-associated microbes. HY provided the rice root samples. LXW and YL wrote the manuscript. LD, MXC, YB, HY, ACH, JF, TS and YGL reviewed and edited the manuscript. All authors approved the final manuscript.

## AVAILABILITY OF DATA AND MATERIALS

The RNA-Seq data and Whole genome sequences of rice-associated isolates generated in this study were deposited to the European Nucleotide Archive (ENA) under the project accession PRJEB70882.

## DECLARATIONS

### Ethics approval and consent to participate

Not applicable.

### CONSENT FOR PUBLICATION

Not applicable.

### DECLARATION OF INTERESTS

The authors declare no competing interests.

## Notes

### Competing Interest Statement

The authors have declared no competing interest.

